# The shared and distinct roles of *Prochlorococcus* and co-occurring heterotrophic bacteria in regulating community dynamics

**DOI:** 10.1101/2025.09.25.678681

**Authors:** Christine A. Ziegler, James I. Mullet, Allison Coe, Nhi N. Vo, Eli Salcedo, Dillon M. Arrigan, Sierra M. Parker, Sallie W. Chisholm

**Affiliations:** Department of Civil and Environmental Engineering, Massachusetts Institute of Technology, Cambridge, MA, USA; Department of Biology, Massachusetts Institute of Technology, Cambridge, MA, USA

## Abstract

*Prochlorococcus* is the world’s most abundant photosynthetic organism with over 10^27^ cells distributed across much of Earth’s oceans, and is collectively responsible for almost 10% of marine carbon fixation. Naturally co-occurring heterotrophic bacteria at roughly 10^5^-10^6^ cells mL^-1^ in the oceans have been shown to increase *Prochlorococcus* fitness and productivity. Despite this massive scale, our understanding of these globally important interactions remains limited, with past research largely focused on single *Prochlorococcus*-heterotroph pairings involving only a few species. In this study, we extend this perspective by using synthetic communities containing multiple diverse heterotrophic strains isolated from *Prochlorococcus* enrichment cultures. Specifically, we isolated the four most abundant co-occurring heterotroph species and examined both individual *Prochlorococcus*–heterotroph interactions and interactions within a synthetic community comprising *Prochlorococcus* and all four heterotrophs. Using absolute quantification of RNA, DNA, and cell counts over the course of *Prochlorococcus* growth curves, we find that *Prochlorococcus* has a modest, species-independent transcriptional response to heterotrophs, whereas each heterotroph displays a markedly different transcriptional response to the community and fulfills distinct metabolic roles. Transcriptional analyses reveal several potential crossfeeding interactions and indicate that community dynamics are influenced not only by metabolic activity but also antagonistic mechanisms, defense responses, and coordinated group behaviors. By pairing synthetic community approaches with absolute abundance measurements, we can gain deeper insight into the forces that shape microbial community assembly in the oceans and their role in driving the global carbon cycle.

## Introduction

*Prochlorococcus* is the world’s most abundant photosynthetic organism, with over 10^27^ cells inhabiting Earth’s oceans collectively contributing to around 10% of marine primary productivity^1,2^. Despite its minimal and streamlined genome (< 2 Mb) encoding for ∼1,700-2,000 genes^3^, *Prochlorococcus* is highly diverse, with strains divided into two ecotypes: high-light (HL) and low-light adapted (LL), based on optimal light conditions for growth^4,5^. HL strains, such as MED4, with the smallest genomes and cell sizes among *Prochlorococcus*, are found throughout the euphotic zone but reach peak abundance in its upper layers^6,7^, where nutrients are more limited and stress from UV radiation is more prominent.

In the upper euphotic zone, ∼10^4^-10^5^ cells mL^-1^ of *Prochlorococcus* co-exist with ∼ 10^5^-10^6^ cells mL^-1^ of heterotrophic bacteria^8,9^ and are often co-isolated because of their similar cell sizes and the isolation techniques employed. Xenic *Prochlorococcus* cultures, which include a diverse assemblage of heterotrophs (ranging from 1 to 25 unique ASVs in cultures analyzed thus far^10^ can be used to isolate these co-occurring heterotrophs. Axenic *Prochlorococcus* cultures (pure cultures that are free of heterotrophs) can then be established by dilution-to-extinction techniques using media containing compounds such as pyruvate, which help mitigate oxidative stress^11–13^, which *Prochlorococcus* is particularly sensitive to as it lacks the catalase gene, *katG*^*13*,14^. Heterotrophs can then be reintroduced to axenic *Prochlorococcus* cultures to examine *Prochlorococcus*-heterotroph interactions, which can range from agnostic to beneficial depending on the *Prochlorococcus*-hetreotroph pairing^15,16^. If the pairing is beneficial, *Prochlorococcus* can exhibit higher growth rates and increased resilience to phage infection, temperature fluctuations, and oxidative stress^13,17-19^. Heterotrophs can even extend *Prochlorococcus’* survival under extreme nutrient starvation^20,21^ and dark stress by influencing its metabolism to utilize metabolites produced by the heterotrophs^22,23^.

While these studies have provided key insights into *Prochlorococcus*-heterotroph interactions, they have been limited by investigating the relationship between *Prochlorococcus* and a single heterotroph, and only a few have studied *Prochlorococcus* in the context of a heterotroph isolated from its native community^13,15,24,18^. Here we build on this progress by isolating the most abundant native heterotrophs from a xenic culture of MED4, a high-light adapted *Prochlorococcus* strain that is abundant at the ocean surface, and constructed both single co-cultures and synthetic communities to address the following key questions: How does *Prochlorococcus* respond to its native heterotrophs? What role does each member play in community fitness? And how do these dynamics differ from their behavior in single co-cultures?

## Results and Discussion

### Development of synthetic co-cultures and community

To develop ecologically-relevant co-cultures and a synthetic community, we first determined which heterotrophic bacteria were present in our xenic *Prochlorococcus* MED4 culture (hereafter referred to as MED4)^25^. Metagenomic sequencing revealed that MED4 accounted for the majority of the culture, while heterotrophs comprised the remaining ∼20% (Fig. 1A). Within this heterotroph fraction, we identified six bacterial families present at >0.5% read abundance We set out to isolate representative strains from as many of these six bacterial families as possible using several recipes of seawater-based agar plates (see Methods). From this effort, we successfully obtained, sequenced, and assembled genomes from one representative each of *Marinobacteraceae* (*Marinobacter*), *Alteromonadaceae* (*Alteromonas*), and *Phyllobacteriaceae* (*Pseudohoeflea*), as well as two strains of *Thalassospiraceae* (*Thalassospira*) (Fig. 1B, Supp. Table 1). These isolated strains represent the four most abundant heterotroph families that together make up ∼90% of the heterotroph population in the MED4 xenic culture (Fig. 1A).

**Figure 1:**
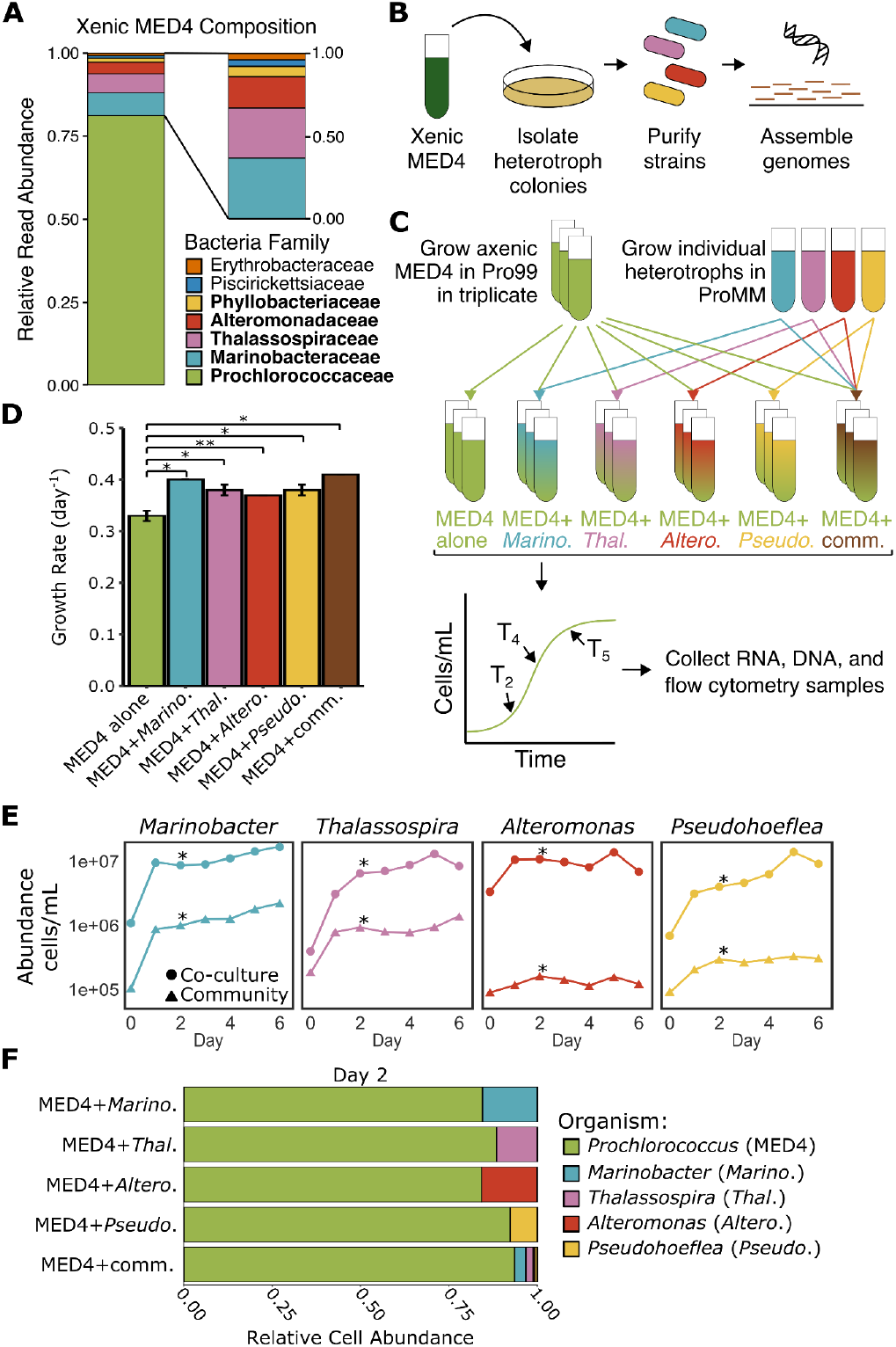
Construction and growth dynamics of synthetic heterotroph communities derived from a xenic *Prochlorococcus* MED4 culture. **A)** Relative read abundances of all bacteria families in MED4 xenic cultures (left) and relative abundances of families within the heterotroph fraction only (right). **B)** Schematic for the isolation and characterization of heterotroph strains from the MED4 xenic culture. **C)** Schematic of the experimental setup for constructing the co-cultures and synthetic community. MED4 cultures and all co-cultures were grown on Pro99 (no added organic carbon), while heterotrophs were grown on ProMM (with added organic carbon and vitamins) prior to being washed and added to co-cultures. Heterotroph abbreviations: Marino=*Marinobacter*, Thal=*Thalassospira*, Altero=*Alteromonas*, Pseudo=*Pseudohoeflea*, and Comm=Community **D)** MED4 growth rates for each of the conditions from **C**. Bars show the mean of three biological replicates, with error bars representing the standard deviation. Asterisks indicated statistical significance: p<0.05 (*), p<0.01 (**). **E)** Growth curves for the four heterotrophs in single co-culture (circles) or in the synthetic community (triangles). Asterisks denote RNA sampling (Day 2). **F)** Relative abundances of bacteria in the co-cultures and synthetic community (Day 2), as measured by flow cytometry and metagenomic read mapping after correction for extraction efficiency. Bars represent the mean of three biological replicates.

### Prochlorococcus growth

Armed with these five heterotrophs and an axenic (pure) MED4 culture, we next established various cultures: MED4 axenic (control), MED4 with each of the isolated heterotrophs (co-cultures), and MED4 with all the isolated heterotrophs (community) in order to determine how MED4 responds to individual heterotrophs and a community of heterotrophs, and how each heterotroph contributes to MED4 and community fitness (Fig. 1C). Using flow cytometry (Supp. Table 2) and metagenomic sequencing (Supp. Table 3), we found that the addition of individual heterotrophs, as well as the community of heterotrophs, significantly increased the growth rate of MED4 (*t-test*, p<0.05; Fig. 1D). Although no studies have examined *Prochlorococcus* growth with isolates of *Thalassospira* or *Pseudohoeflea*, a study with a different *Marinobacter* isolate demonstrated no measurable effect on MED4 growth rate^15^. Studies of *Prochlorococcus* with *Alteromonas* have yielded conflicting results: one showed a slight increase in *Prochlorococcus* growth rate^18^, while another found no significant difference in growth rate^24^. These discrepancies likely reflect differences in *Prochlorococcus* and heterotroph strains, *Prochlorococcus*:heterotroph inoculum ratios, and experimental conditions. In our experiments, MED4 displayed slight, but statistically significant increases in growth rate when co-cultured with any of our four co-isolated heterotrophs or in the community of these heterotrophs, highlighting how naturally co-occurring heterotrophs can influence MED4 fitness (Fig. 1D). Given *Prochlorococcus*’ abundance in the oceans, even small growth rate changes could have a large impact on marine carbon cycling.

### Heterotroph growth

As previously mentioned, we independently isolated two *Thalassospira* strains. After genome assembly and sequencing, we noticed that the two *Thalassospira* strains had nearly identical genomes, with only four single nucleotide variants (SNVs) and a putative 2Mb inversion (Supp. Fig. 1, Supp. Table 4). To test whether these genetic differences contributed to phenotypic ones, both strains were included in all individual co-cultures and the synthetic community, effectively doubling the total *Thalassospira* abundance relative to the other heterotroph strains in the community culture. We did not detect any significant phenotypic differences in the single co-cultures, either from MED4 or *Thalassospira* gene expression (Supp. Fig. 2, Supp. Table 5); therefore, for simplicity, only one strain of *Thalassospira* is discussed here on..

Given that the presence of heterotrophs increased the growth rate of MED4 (Fig 1D), we next asked how each heterotroph’s growth was affected by co-culture with MED4 versus within the synthetic community. Using flow cytometry (Supp. Table 2), we found that in all co-cultures with MED4, heterotrophs exhibited an initial growth spike within the first 24 hours, similar to that observed by Becker *et al*^16^. Following this spike, *Alteromonas* plateaued, while *Marinobacter, Thalassospira*, and *Pseudohoeflea* continued to grow more gradually (Fig. 1E), suggesting that all heterotrophs initially took up MED4 photosynthate but then slowed down their growth rate to match the rate of organic carbon production by MED4. In the synthetic community, using both flow cytometry and metagenomic sequencing (Supp. Table 3), we observed similar growth patterns, where *Marinobacter, Thalassospira*, and *Pseudohoeflea* grew slowly, and *Alteromonas* plateaued (Fig. 1E). Overall, heterotroph growth trends were comparable between single heterotroph co-cultures with MED4 and the full community, although absolute abundances of heterotrophs were lower in the community due to smaller initial inocula of each strain, as all cultures were inoculated with the same total number of heterotrophs. These similar growth patterns suggested that the presence of other community members did not drastically affect their individual growth phenotypes. Throughout the MED4 growth curves, each heterotroph comprised only a small fraction of the total cells, with MED4 comprising 80-90% of the total cells and *Marinobacter* and *Thalassospira* dominating the heterotroph pool (Fig.1F, Supp. Fig. 3). This relative ratio of *MED4:*heterotrophs mirrored that in our xenic cultures that were isolated from the wild (Fig. 1A), suggesting our synthetic communities recapitulate natural dynamics and provide a useful tool for investigating the mechanisms underlying community interactions.

### Transcriptional Response of Prochlorococcus to the presence of heterotrophs

We next wondered whether the increase in MED4 growth rate observed in the presence of individual heterotrophs and the synthetic community (Fig. 1D) was due to a common mechanism and whether MED4 transcription profiles could provide mechanistic insight. To this end, we performed RNA-sequencing on cultures in mid-exponential growth (Day 2, ∼48 hours after inoculation). MED4 co-cultures with *Marinobacter, Thalassospira*, and *Pseudohoeflea* had the most significantly differentially expressed MED4 genes when compared to growth of axenic MED4 (Fig. 2A): 1,077/1,921 genes, 1,049/1,921 genes, and 1,221/1,921 genes, respectively (Supp. Table 6). In contrast, MED4+*Alteromonas* exhibited few significant changes (63/1,921), likely due to a larger variance among biological replicates (Supp. Fig. 4). Notably, we did not detect any significantly differentially expressed genes in the MED4+community condition, possibly due to the competitive nature of a community, where some heterotrophs may consume metabolites produced by the others before MED4 is able to.

**Figure 2:**
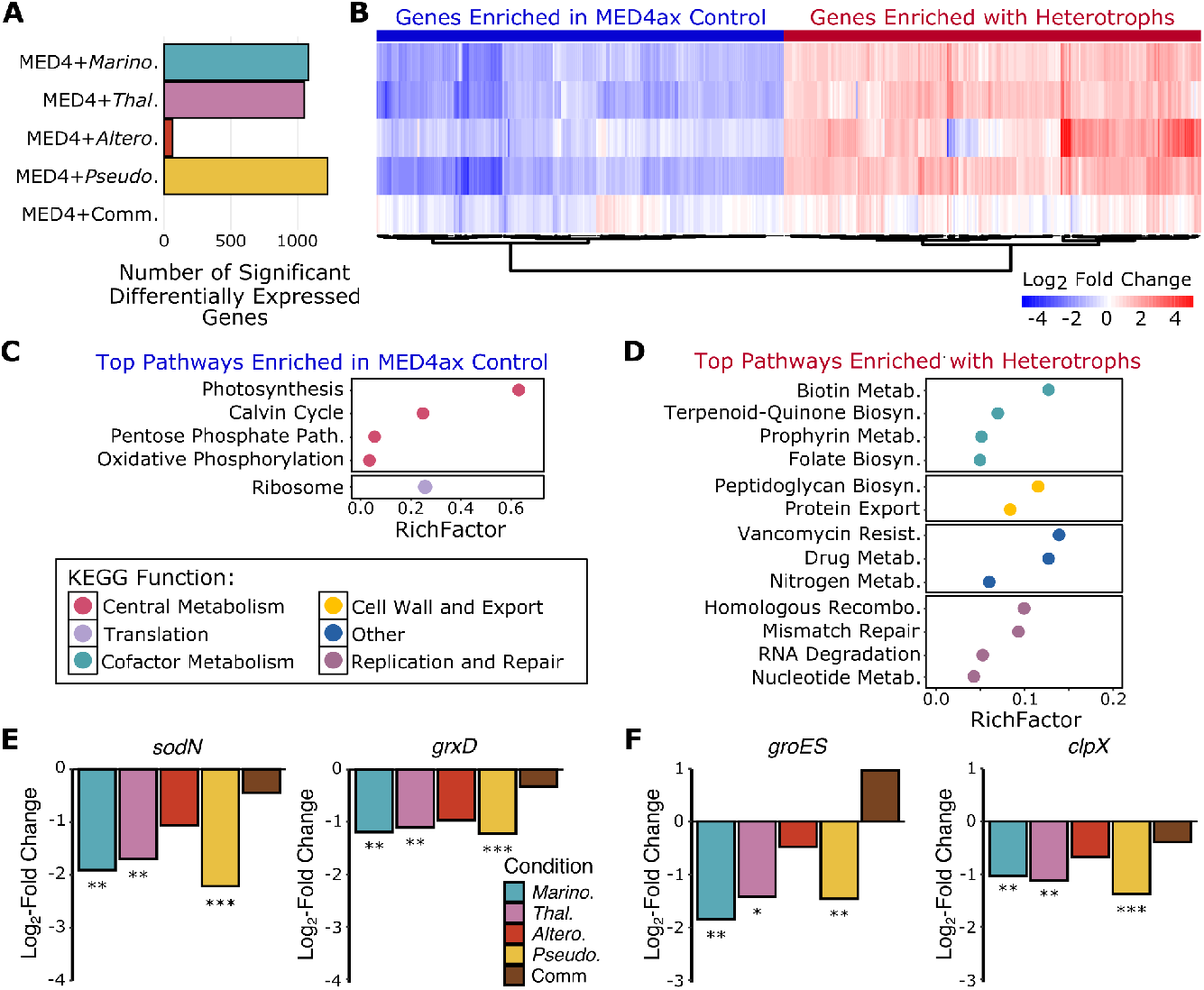
Transcriptional response of MED4 to the presence of heterotrophs and their effects on oxidative stress and protein damage. **A)** Number of significantly differentially expressed MED4 genes in each condition relative to axenic MED4 . Heterotroph abbreviations: Marino=*Marinobacter*, Thal=*Thalassospira*, Altero=*Alteromonas*, Pseudo=*Pseudohoeflea*, and Comm=Community. Significance is defined as a log_2_ fold change > +/- 1.2 and q<0.05. **B)** Heatmap of the log_2_ fold changes for all MED4 genes significantly differentially expressed in at least one condition relative to growth alone. Row labels correspond to panel **A** labels. Genes are clustered on the x-axis by similarity across treatments. Negative values in blue indicate higher expression in axenic MED4, and positive (red) values indicate higher expression in MED4 grown with heterotrophs.**C)** Top pathways enriched among genes more highly expressed in MED4 grown alone relative to growth with heterotrophs, colored by KEGG function. **D)** Top pathways enriched among genes more highly expressed in MED4 grown with heterotrophs relative to growth alone. **E)** Representative MED4 oxidative stress genes and their log_2_ fold change in co-culture relative to growth alone. Asterisk indicate statistical significance q<0.05 (*), q<0.01(**), and q<0.001(***). **F)** Same as **E**, but for representative protein damage-related genes.

Surprisingly, MED4 exhibited a very similar transcriptional response to each of the four heterotrophs relative to growth alone, suggesting a generalized, species-agnostic response to heterotrophs (Fig. 2B). This is striking given the diverse phenotypic and metabolic capabilities of the heterotrophs (as described in more detail below). Most MED4 genes significantly differentially expressed with one heterotroph were also significantly differentially expressed with at least one other (Supp. Fig. 5, Supp. Table 6), and even genes that did not reach statistical significance showed similar expression trends (Fig. 2B).

To elucidate MED4’s generalized transcriptional response to heterotrophs, we next assessed the top pathways with decreased or increased expression in MED4 co-cultures relative to axenic MED4 (Fig. 2C,D) In axenic cultures (relative to co-cultures), MED4 showed the strongest increased expression in pathways related to photosynthesis, ribosome biosynthesis, oxidative phosphorylation, and the pentose phosphate pathway (Fig. 2C). When grown axenically, MED4 relied solely on carbon fixation via photosynthesis, a process that generates high oxidative stress^26,27^ and likely necessitates continuous replacement of damaged proteins. This is supported by the higher expression of redox stress genes (Fig. 2E) and protein damage-related genes (Fig. 2F) in axenic cultures compared with co-cultures. In contrast, co-culture with heterotrophs (relative to axenic growth) led to increased expression of MED4 pathways involved in B vitamin and cofactor biosynthesis, cell wall biosynthesis and protein export, antimicrobial resistance, nitrogen metabolism, and replication and repair (Fig. 2D). Similar transcriptional patterns have been reported in MED4*-Alteromonas* co-cultures^18,28^. Freed from the need to devote energy primarily to photosynthesis and protein turnover / repair, MED4 can redirect energy towards metabolizing heterotroph-derived compounds while experiencing reduced oxidative stress. These results not only replicated previous findings in MED4-*Alteromonas* co-cultures^18,28^ despite different experimental designs and *Alteromonas* strains, but also demonstrated that MED4’s transcriptional response to its most abundant native heterotrophs is generalized and universal.

### Transcriptional response of individual heterotrophs to growth in the community

Having characterized MED4’s transcriptional response to heterotrophs, we next asked how each heterotroph’s transcriptome is affected by growth with MED4 alone versus the full synthetic community. After correcting for library size and normalizing to internal standards (Supp. Table 7), each heterotroph displayed distinct transcriptional responses in the community relative to when individually co-cultured with MED4 (Fig. 3A, Supp. Table 8). *Marinobacter*, which grew well under both conditions, had few transcriptional differences in major metabolic pathways, except for genes involved in energy metabolism and translation, which were increased in expression in the community (Fig. 3A). This likely explains its slightly faster growth rate in the community at this stage of the MED4 growth curve (Fig. 1E) and suggests that *Marinobacter* is the more dominant heterotroph. Although *Alteromonas* remained mostly in stationary phase under both culturing conditions, genes involved in energy, vitamin, nucleotide, carbohydrate, lipid metabolism, and terpenoid biosynthesis had slightly higher expression in the community (Fig. 3A), suggesting a more active metabolic state and contribution of metabolites to the community despite limited growth (Fig. 1E).

**Figure 3:**
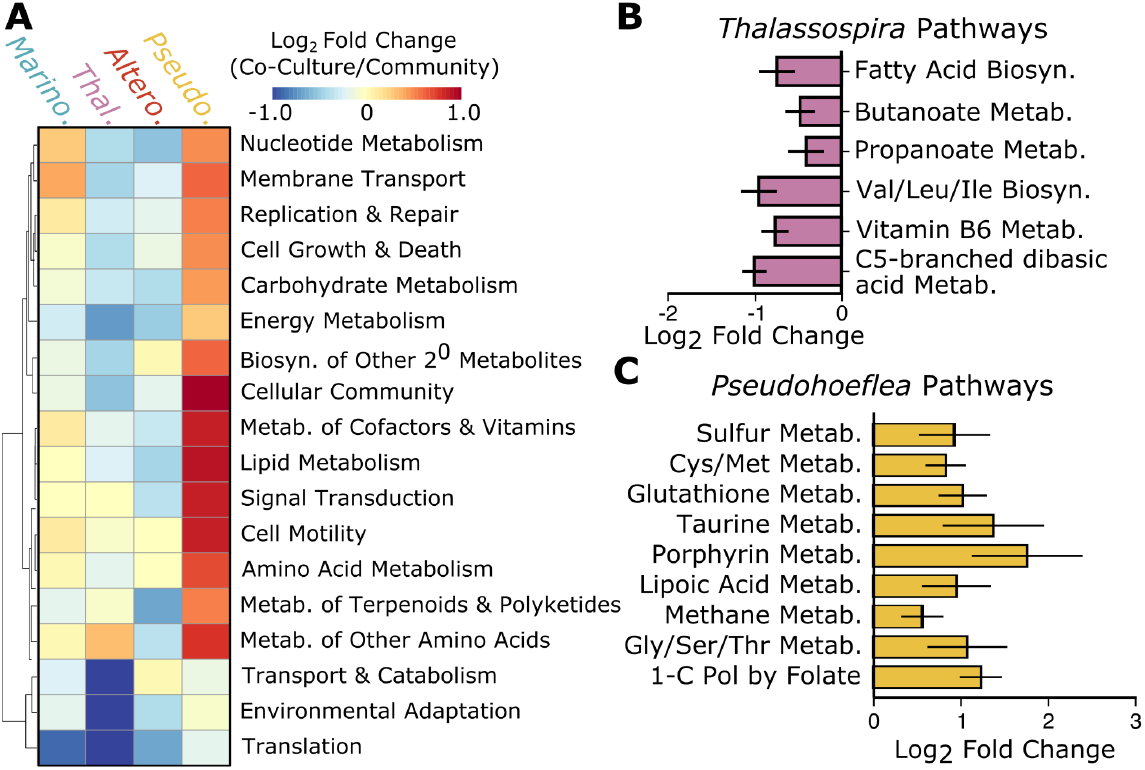
Transcriptional and metabolic responses of each heterotroph to growth in the synthetic MED4 community relative to growth in co-culture with MED4. **A)** Heatmap of the mean log_2_ fold change in transcript abundance for each major KEGG pathway for each heterotroph when growth in individual co-culture versus its growth in the synthetic community. Positive (red) values indicate higher expression in the co-culture relative to the community. **B)** Mean log_2_ fold change in transcript abundance for pathways involved in aliphatic carbon skeleton metabolism in *Thalassospira* co-culture versus community. **C)** Same as **B**, except for pathways involved in sulfur and one-carbon metabolism in *Pseudohoeflea*.

*Thalassospira* had even more pronounced increased expression in numerous pathways in the community relative to growth in the co-culture, including translation, vitamin metabolism, transport and catabolism, and energy, nucleotide, carbohydrate, lipid, and amino acid metabolism (Fig. 3A). Almost all of the pathways for aliphatic carbon metabolism were among the top 15 sub-pathways with increased expression, including fatty acid biosynthesis, butanoate and propanoate metabolism, vitamin B6 metabolism (an essential cofactor used in fatty acid and amino acid biosynthesis), and C5-branched dibasic acid and branched-chain amino acid (leucine, isoleucine, and valine) (Fig. 3B). These findings suggest that other organisms, likely *Marinobacter* due to its relatively unchanged transcriptome, consumed MED4-derived carbon skeletons in the community, requiring *Thalassospira* to synthesize more of its own lipids and branched chain amino acids to support its growth in the community. In contrast, most metabolic pathways had decreased expression in *Pseudohoeflea* in the community, with the top pathways with decreased expression primarily involved in sulfur and one-carbon metabolism, which are tightly linked through cysteine and methionine metabolism (Fig. 3C). This may indicate either reduced availability of sulfur and one-carbon metabolites available to *Pseudohoeflea* in the community, or increased synthesis of cysteine and methionine in co-culture as defense against oxidative stress from MED4, as studies in *E. coli* have revealed that these metabolites can serve as a cellular defenses against reactive oxygen species^29^. Overall, these results highlight that while MED4 exhibits a similar transcriptional response to heterotrophs, each heterotroph has a unique transcriptional and metabolic response to the community. MED4, as the base of the food web, maintains steady organic carbon production, whereas the heterotrophic community drives the dynamic metabolism and collective carbon processing.

### Cross-feeding within the community

Given the notably different transcriptional responses of the heterotrophs to the community relative to single co-culture, we next wondered what metabolic roles each organism played within the community. Using cell counts and internal standard-corrected transcript counts to determine absolute abundance (Supp. Table 9), we first quantified the fraction of transcripts derived from each organism within the community for each gene and metabolic pathway. We note that transcript abundance is not a direct measurement of metabolic activity, but it is a common proxy due to its correlation in bacteria^30^. Overall, MED4 expressed the vast majority of transcripts for lipid, nucleotide, and amino acid metabolism (Fig. 4A), while *Marinobacter* performed most of the carbohydrate and energy metabolism. Although this distribution reflects their numerical abundance in the community, it was notable how few transcripts from each of the remaining three organisms contributed to these major macromolecule metabolic processes.

**Figure 4:**
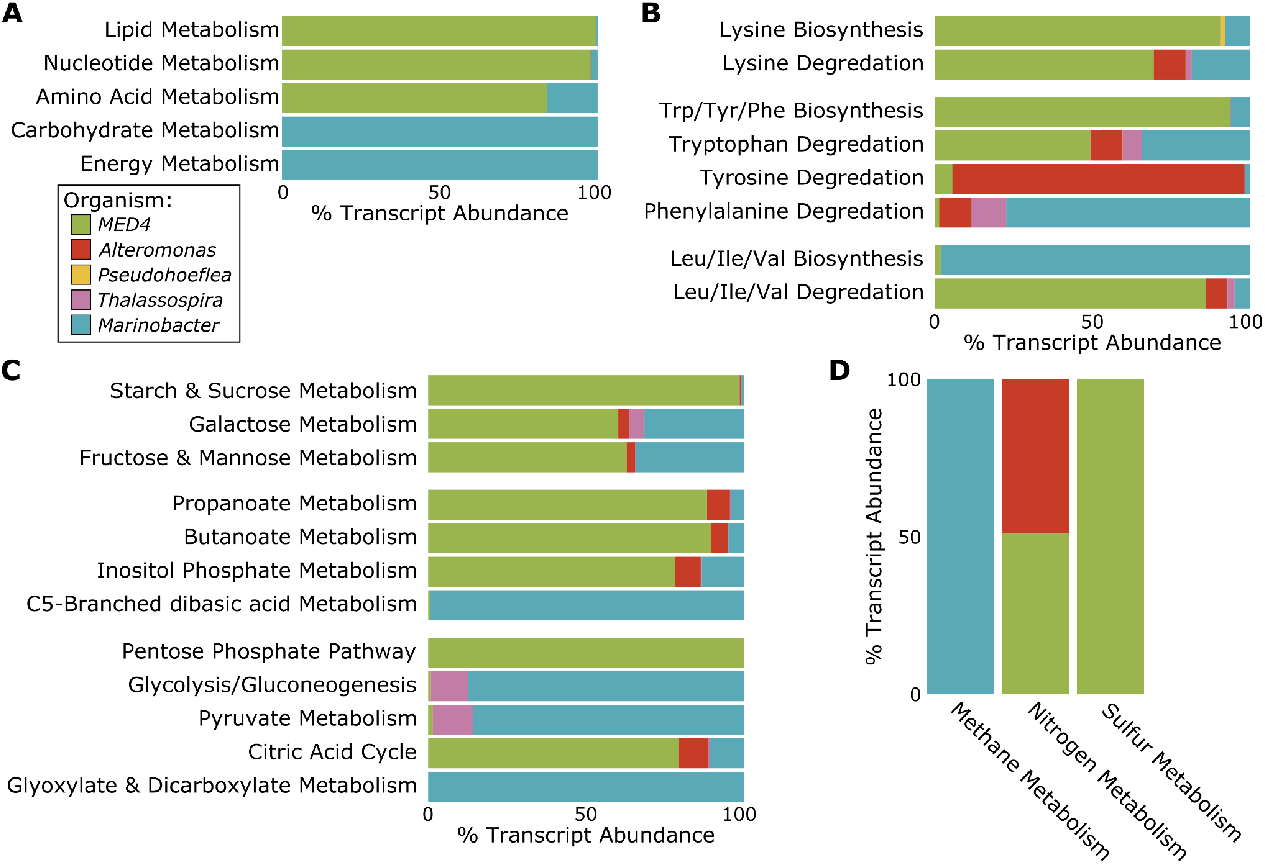
Crossfeeding in the synthetic community. **A)** Mean percentage of total transcripts contributed by each community member for each core gene within major macromolecular metabolic pathways in the synthetic community. Colors represent different organisms. **B)** Examples of biosynthetic and degradation pathways for amino acids. **C)** Examples of carbohydrate and energy metabolic pathways. **D)** Metabolic pathways for one-carbon units, nitrogen, and sulfur.

Since MED4 only contributed ∼80% of the transcripts for amino acid metabolism (Fig. 4A), we asked whether different heterotrophs contributed to particular amino acid metabolic pathways, and if this could provide clues into cross-feeding interactions. As was expected given its role in organic carbon production, most amino acid synthesis and degradation appeared to be driven by MED4. However, transcript profiles of genes for the metabolic pathways of lysine, aromatic amino acids, and branched-chain amino acids displayed patterns indicative of cross-feeding (Fig. 4B). The expression patterns imply that *Marinobacter* and *Alteromonas* consumed MED4-derived lysine in the community. Similarly, *Marinobacter, Alteromonas*, and to a lesser extent, *Thalassospira* consumed MED4-derived aromatic amino acids (tyrosine, phenylalanine, and tryptophan), consistent with metabolic profiles observed in *Prochlorococcus* spent-media^31^. Conversely, our absolute transcript abundance data suggest that the branched-chain amino acids (leucine, isoleucine, and valine) were synthesized almost entirely by *Marinobacter*, which MED4 and, to a small extent, *Alteromonas*, were able to consume. Recent studies revealed that MED4 is able to consume organic carbon from heterotrophic bacteria^22,23,28^, but the species of organic carbon and specific cross-feeding interactions were mostly uncharacterized. These lysine, aromatic amino acid, and branched-chain amino acid metabolism findings suggest a few possible routes for cross-feeding of organic carbon species among MED4 and community members.

We next examined whether any similar patterns of cross-feeding would be evident in sub-pathways of carbohydrate and energy metabolism since these major pathways were largely dominated by *Marinobacter*, but are essential for life in all organisms. In the wild, heterotrophic bacteria generally derive energy from either sugars or organic acids^32^, both of which get funneled into the Krebs (citric acid) cycle either directly (acids) or through glycolysis and pyruvate (sugars). Transcript abundances in the community suggest that MED4 synthesized starch and sucrose, some of which *Marinobacter* consumed (e.g., fructose, galactose, and mannose) and funneled into glycolysis and pyruvate metabolism (Fig. 4C). *Thalassospira* also utilized galactose to generate pyruvate, whereas MED4 and *Alteromonas* primarily relied on short chain fatty acids and lipids (butanoate, propanoate, inositol phosphate, and C5-branched dibasic acids) as their primary energy sources (Fig. 4C). Reflective of its numerical dominance in the community, MED4 performed the majority of the Krebs (citric acid) cycle, with some contributions from *Marinobacter* sugar breakdown and *Alteromonas* short chain fatty acid metabolism. Within the Krubs cycle, MED4 produced excess glyoxylate, which was consumed by *Marinobacter* and funneled back into the Krebs cycle. These findings reflect how MED4 and its associated heterotrophs are able to occupy metabolic niches in order to grow in the community.

Finally, we examined transcriptional patterns in the community to assess which organisms were contributing to the utilization and metabolism of carbon, nitrogen, and sulfur, the latter two being replete in the media. *Marinobacter* performed most of the one-carbon metabolism, while MED4 dominated sulfur metabolism and contributed to about half of nitrogen metabolism. *Alteromonas* comprised the other half of nitrogen metabolism transcripts, consistent with its role in lysine uptake and degradation. MED4’s dominance in sulfur metabolism likely reflects its unique membrane, which is composed of sulfolipids instead of canonical phospholipids.

### Roles of MED4 and heterotrophs within specific, essential metabolic pathways

The previous section describes key, large-scale metabolic interactions between community members, but we also wondered whether cross-feeding could occur on even finer-grain scales, such as portions of a metabolic pathway or even just a single enzymatic step. Indeed, this was the case for several of the B vitamin/cofactor biosynthetic pathways. B vitamins serve as essential enzyme cofactors, expanding the repertoire of chemistries cells can perform beyond the functional groups provided by the 20 proteinogenic amino acids. Although cells only require minute amounts of each B vitamin (typically in the nanomolar range found in natural seawater^33^), the transcription, translation, and metabolic investment required to synthesize these molecules is energetically costly^34^.

One such B vitamin, folate (Vitamin B9) had transcripts for its biosynthetic pathway produced almost entirely by heterotrophs in the community and in co-cultures, particularly the later steps in the pathway (Fig. 5A), but the transcripts for folate-utilizing genes were predominantly expressed by MED4. Folate is required for one-carbon metabolism and plays major roles in nucleic acid and protein synthesis, amino acid biosynthesis, and cell division^35^. These results suggest that heterotrophs could enhance MED4 fitness by supplying folate, which MED4 can utilize to support growth and division. The transcripts for the biosynthetic pathway of NAD^+^, generated from Vitamin B3, are also produced in a similar manner (i.e. mainly by the heterotrophs) in the community from quinone (Supp. Fig. 6A). NAD^+^ can be phosphorylated into NADP^+^ by MED4 and serves as a powerful antioxidant, yet another means by which heterotrophic bacteria could help mitigate oxidative stress in MED4 (Supp. Fig. 6B-C).

**Figure 5:**
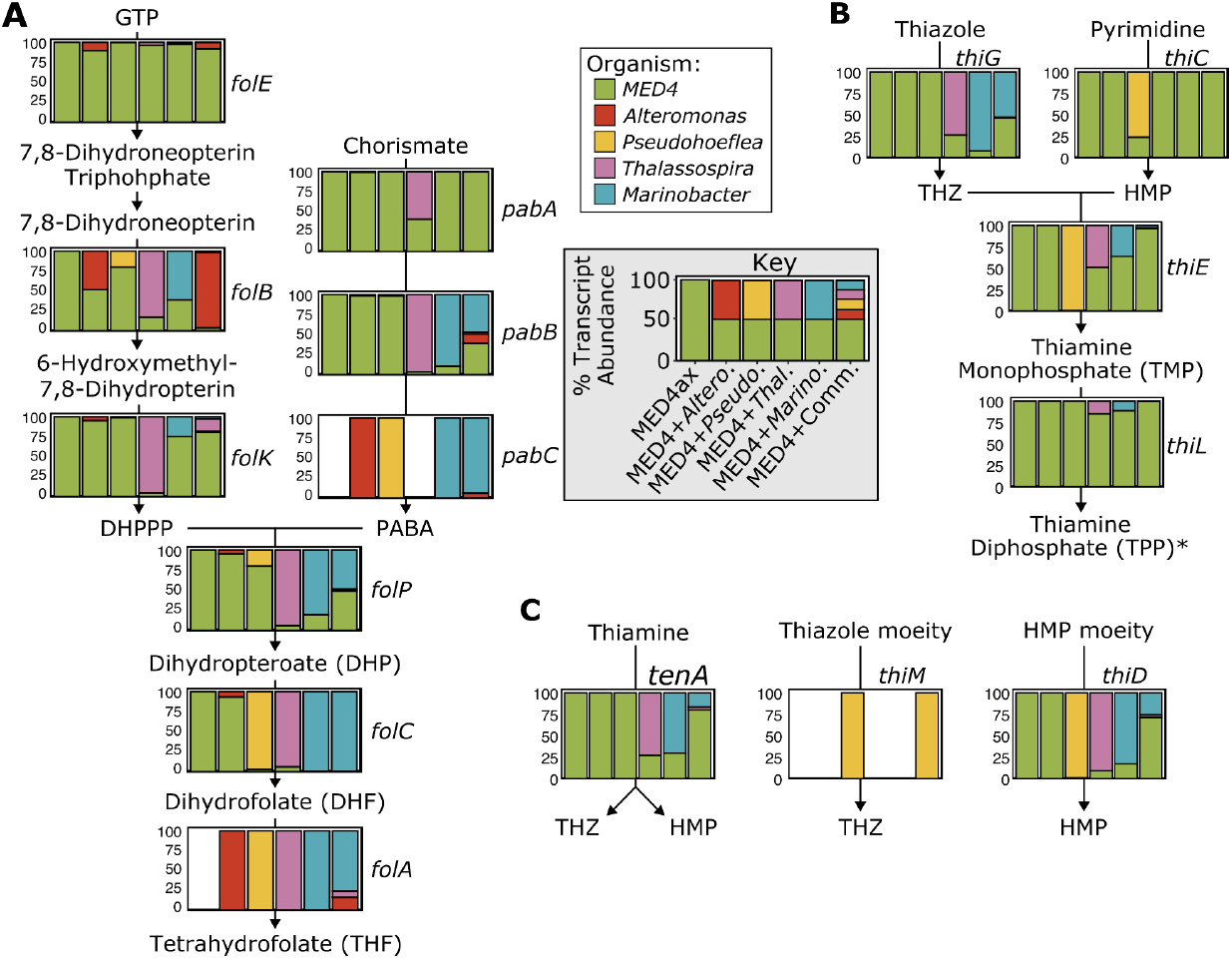
Transcript contribution of each organism to each step of two B vitamin biosynthetic pathways. Mean transcript abundances (%) for MED4 (green), *Alteromonas* (red), *Pseudohoeflea* (yellow), *Thalassospira* (purple), and *Marinobacter* (blue) are shown for each gene in the following pathways: **A)** Folate (Vitamin B9) biosynthesis, **B)** Thiamine (Vitamin B1) biosynthesis (*Pseudohoeflea* does not express *thiG*, and only MED4 and *Pseudohoeflea* express *thiC*), and **C)** Thiamine salvage.See the “Key” in the gray box for x-axis labels. Absence of a colored bar indicates that no organism in that condition encodes that gene. Abbreviations for metabolites as follows: THZ=thiazole precursor, HMP=4-amino-5-hydroxymethyl-2-methylpyrimidine.

While the two B vitamin biosynthetic pathways discussed above served as examples of heterotrophs likely producing compounds beneficial to MED4 by generating the majority of transcripts for performing the final few metabolic steps of the pathways, we found that MED4 could provide heterotrophs with vitamin B1, through expression of a single gene, either *thiC* (*Alteromonas, Marinobacter*, and *Thalassospira*) or *thiG* (*Pseudohoeflea*) (Fig. 5B). Thiamine pyrophosphate (TPP) is required for several steps in the Krebs cycle and is therefore essential for energy production and life. TPP biosynthesis requires two major precursors: a thiazole precursor (THZ, produced by *thiG*) and 4-Amino-5-hydroxymethyl-2-methylpyrimidine (HMP, produced by *thiC*). Many marine heterotrophs have been reported to lack *thiC* and are therefore auxotrophic for HMP^36,37^. Instead, they rely on salvage mechanisms, such as *tenA* and *thiD* to acquire HMP from their environment (Fig. 5C). While all four heterotrophs used in this study contain all the genes required for thiamine biosynthesis (Supp. Fig. 7), only *Pseudohoeflea* expressed *thiC* and could produce its own HMP. *Marinobacter, Thalassospira*, and *Alteromonas* instead appear to acquire HMP from the *thiD* salvage pathway (Fig. 5C). *Pseudohoeflea*, however, did not express *thiG* to produce THZ and instead relied on *thiM* to acquire this component via the salvage pathway (Fig. 5C). Unlike the four heterotrophs, MED4 expressed every essential step in thiamine biosynthesis and thus could supply thiamine or its metabolic precursors to the heterotrophic community. To our knowledge, this is the first time *Prochlorococcus* has been shown to provide essential metabolites to the surrounding community of microorganisms aside from organic carbon.

### Behaviors of each organism within the community

While both the coarse- and fine-scale metabolic and cross-feeding examples described above provide some insight into community dynamics and niche partitioning, they do not fully explain the growth patterns and relative abundance of each organism in the community. To tackle this question, we assessed the relative transcript abundance of the major cell-cell communication pathways (Fig. 6A). Interestingly, many of the community members expressed genes for the biosynthesis of antibiotics (Fig. 6B). While it is likely they do not produce lethal amounts since all strains grew in our experiments, it is possible antibiotics are used as a means of reducing competition for resources. MED4, *Marinobacter* and *Alteromonas* produce the most antibiotic biosynthesis transcripts, and MED4, *Marinobacter*, and *Thalassospira* expressed the most antibiotic resistance transcripts (Fig. 6C). This resonates with the relative growth patterns observed in the community cultures, where MED4, *Marinobacter*, and *Thalassospira* (all expressing the majority of antibiotic resistance genes) grew the best in the community followed by *Pseudohoeflea*, then *Alteromonas*.

**Figure 6:**
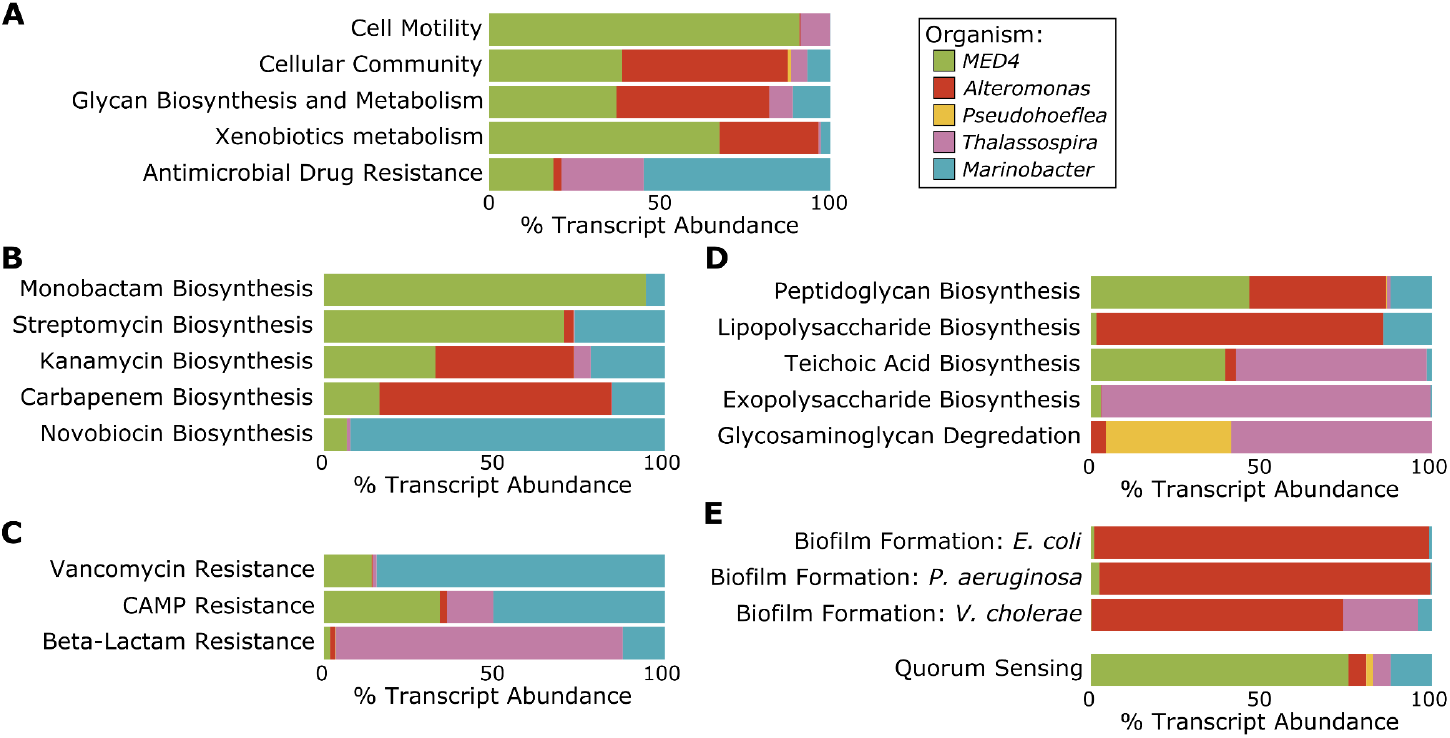
Distinct behaviors of each organism within the community. Mean percentage of transcripts each organism contributes in the synthetic community to the following pathways: **A)** cell-cell communication, **B)** antibiotic biosynthesis, **C)** antibiotic resistance, **D)** cell wall and exopolysaccharide biosynthesis and degradation, **E)** biofilm and quorum sensing,.

Based on transcript profiles, we hypothesize that *Marinobacter* produced antibiotics and respective resistance genes to outcompete other heterotrophs and thereby gained preferential access to MED4-derived organic compounds. *Thalassospira* expressed the majority of transcripts for cellular motility via chemotaxis and flagellae pathways (Fig. 6A) and was able to survive by traveling towards nutrients while evading attacks by antibiotics via expression of antibiotic resistance genes and generating teichoic acid and exopolysaccharide to form a protective barrier around the cell (Fig. 6D)^38,39^. *Alteromonas*, meanwhile, produced a large portion of peptidoglycan and lipopolysaccharide (Fig. 6D), which were used to construct a strong, protective cell wall^40,41^, and dominated the transcript pool for biofilm formation genes (Fig. 6E). While metabolically active, *Alteromonas* did not grow as quickly in the community and was likely in a biofilm, which served as a protective barrier from the antibiotics it produces for others. *Pseudohoeflea* was able to survive by not being metabolically active and, therefore, was not as sensitive to antibiotic stressors. From this analysis, it is clear that each organism has a distinct role in the community. However, these community roles were not observed in individual co-cultures, highlighting the importance of studying communities rather than just single interactions.

### *MED4* and heterotroph transcriptional responses over the growth curve

All transcriptomic responses discussed above correspond to Day 2 (∼48 hours after inoculation), representing the mid-exponential phase of the MED4 growth curve. We next asked how the community members would acclimate and interact at later stages of the growth curve, specifically Day 4 (late-exponential growth phase) and Day 5 (transition point between late-exponential and stationary phase). Overall, relative transcriptional responses of both MED4 and the heterotrophs to each other and the community diminished over time, with Day 2 consistently showing the strongest differential expression (log_2_ fold change). The magnitude of differential expression was reduced on Day 4 and further diminished by Day 5 (Supp. Fig. 8, Supp. Fig. 9).

MED4 expression patterns on Day 4 closely resembled those from Day 2 (Supp. Fig. 8), but by Day 5, no genes met the criteria for significant differential expression (log_2_ fold change +/- 1.2, q-value ≤ 0.05). This could reflect increased transcriptional variability as cultures transition into stationary phase, or suggest that heterotrophs enhance MED4 fitness primarily through early transcriptional effects. While the overall transcriptional changes in the heterotrophs also decreased over time, subtle shifts in the top enriched pathways were still observed (Supp. Fig. 9), suggesting ongoing acclimation to community context even as cultures approached stationary phase.

## Conclusion

Our exploration of the collective metabolism of individual MED4-heterotroph co-cultures and the synthetic community has revealed that overall, MED4 has a modest, generalized transcriptional response to the presence of its naturally co-occurring heterotrophs, at least in part due to the heterotrophs’ mitigation of the oxidative damage caused by photosynthesis. Each heterotroph, on the other hand, possesses different metabolic capabilities and plays a distinct role in the community (Fig. 7). Within the synthetic community, there are many examples of putative cross-feeding with amino acids, sugars, fatty acids, and essential B vitamins, indicating that heterotrophs can improve MED4 fitness by supplying it with organic carbon and essential molecules in addition to mitigating oxidative stress.

**Figure 7:**
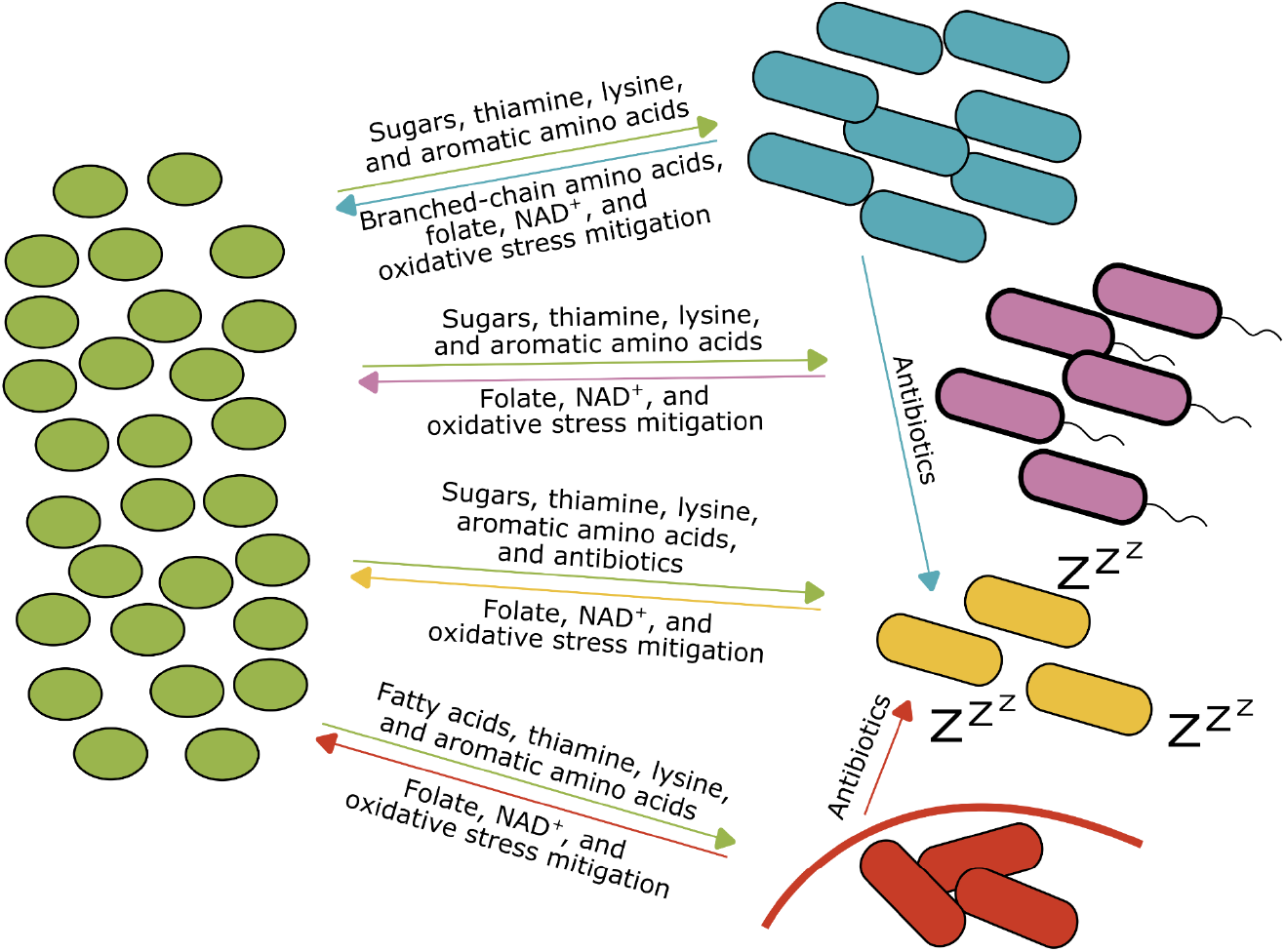
Representative diagram of community interactions. Schematic of interactions between MED4 (green), *Marinobacter* (blue), *Thalassospira* (purple), *Pseudohoeflea* (yellow), and *Alteromonas* (red) within the synthetic community. Arrows next to metabolites are colored by the organism of origin. Cell numbers are an approximation of those found in the synthetic community. Marinobacter is the dominant heterotroph, *Thalassospira* has a thicker cell wall and flagellae, *Pseudohoeflea* is likely not highly metabolically active, and *Alteromonas* forms a protective biofilm, as shown by the red curved line around the cells.

Collectively, the heterotrophs increase the growth rate of MED4 by generating branched-chain amino acids and folate, which is utilized predominantly by MED4 for cell growth and division, as well as NAD^+^, catalase, and superoxide dismutase to help MED4 combat oxidative stress. In addition to supplying the heterotrophs with organic carbon in the form of sugars, aromatic amino acids, lysine, and fatty acids, MED4 also supplies them with thiamine (vitamin B1) and its metabolic precursors, which they do not synthesize on their own. Each heterotroph displayed a unique behavior in the community, with *Marinobacter* dominating the heterotroph population by exuding antibiotics to reduce growth of competitors and acquire MED4-derived metabolites, *Thalassospira* growing well by expressing antibiotic resistance genes, generating strong cell walls, and migrating towards nutrients through chemotaxis and expression of flagellar genes, *Alteromonas* producing antibiotics and forming a protective biofilm with thick cell walls to evade attacks from antibiotics, and *Pseudohoeflea* relying on MED4 and heterotrophs to provide essential nutrients, while reducing its metabolic activity to mitigate the effects of antibiotics.

These behaviors differ from their phenotypes in individual MED4-heterotroph co-cultures, demonstrating the need to study organisms in the context of the community.

Furthermore, the strongest transcriptional responses were observed in mid-exponential phase of the MED4 growth curve, with changes in the MED4 and heterotroph transcriptomes generally decreasing over time. While the findings reported here represent acclimation, rather than steady-state interactions, they are useful for understanding key metabolic processes, cross-feeding, antagonism, and individual phenotypes underlying community formation and dynamics. Overall, our synthetic community approach paired with absolute quantification is a step toward better understanding how individual community members contribute to group fitness in marine microbial assemblages – the complexity of which in the wild is daunting. Because the members of the communities used in our study are self-selected during the original isolation of MED4 – the primary producer upon which they depend – our experimental design represents a natural bridge between studies in the wild and studies with artificial synthetic communities designed from isolates that were not sympatric.

## Acknowledgements

We would like to thank the BPF Genomics Core Facility at Harvard Medical School and the BioMicro Center at MIT for their sequencing expertise and instrument availability that supported this work. We would also like to thank the members of the Chisholm lab for their feedback and suggestions on this manuscript. This work was funded in part by the Simons Foundation Life Sciences Award ID: 337262 and SCOPE Award ID: 721246 to S.W.C.. C.A.Z. was supported by the Simons Postdoctoral Fellowship in Marine Microbial Ecology (LS-FMME-00003951).

## Author Contributions

C.A.Z., A.C., and S.W.C. designed this study. C.A.Z., A.C., E.S., S.M.P., D.M.A., and J.I.M. performed wet-lab experiments and processed samples. J.I.M. and N.N.V. processed the raw sequencing data. N.N.V. normalized RNA-seq reads to the internal standards and assembled the genomes of the heterotrophs. J.I.M. performed the bioinformatic analyses on the resulting data with input from C.A.Z.. C.A.Z. wrote the manuscript with input from all authors.

## Materials and Methods

### Strains and culturing

The xenic *Prochlorococcus* MED4 culture, which was isolated from 5m depth in the Mediterranean Sea in 1989^25^, has been clonally transferred with its co-occurring heterotrophic community for the past 36 years. Cultures were maintained in liquid Pro99 media^42^ in continuous light at 10±1 μmol photons m^-2^s^-1^ in 24°C.

To isolate co-occurring heterotrophs from xenic MED4 for this study, cultures were streaked on ProAC and ProMM agar plates^13,43^. Individual colonies were picked and serially restreaked onto fresh agar plates three times to ensure clonal isolation. Colonies from the final streak were transferred into their respective liquid medium, and DNA was extracted using the DNeasy Blood and Tissue Kit (cat# 69504, Qiagen, Venlo, Netherlands) for 16S Sanger sequencing using universal bacterial 16S primers (8F, 1492R) to confirm identity. Cultures were then cryopreserved in 10% glycerol, flash-frozen in liquid nitrogen, and stored at -80°C. Prior to experiments, cells from freezer stocks were resuspended in 3mL liquid ProMM media and serially diluted to maintain exponential growth until inoculation. On the day of the experiment, heterotroph cells were washed three times to remove exogenous carbon by centrifugation and resuspended in Pro99 media. Cells were enumerated via flow cytometry prior to experimental inoculation.

Axenic *Prochlorococcus* MED4 cultures were grown in liquid Pro99 media^42^ and maintained in balanced growth for at least three transfers in continuous light at 18μmol photons m^-2^s^-1^ in 24°C (growth rates-0.3636, 0.3297, 0.348). Purity of the culture was determined by flow cytometry. On the day of the experiment, biological triplicate cultures were inoculated with axenic MED4 at a density of 2.5e7 cells/mL per treatment and heterotrophs at a total density of 1e6 cells/mL. Synthetic community samples were inoculated with equal cell counts of each heterotroph, altogether equalling 1e6 cells/mL. 225mL cultures were grown in clear polycarbonate 250 mL bottles (Nalgene, Waltham, MA).

### Flow cytometry

Samples were fixed with glutaraldehyde to a final concentration of 0.025%, incubated in the dark for ten minutes, then flash frozen in liquid nitrogen and stored at -80°C until analysis.

Flow cytometry samples were thawed on ice and stained with Sybr Green I (Lonza, Basel, Switzerland) for 55 minutes to analyze DNA content. Samples were analyzed on a Guava 12HT flow cytometer (Cytek Biosciences, Fremont, CA) with excitation using a 488nm blue laser and emissions measured for chlorophyll, cell size, and SYBR-stained DNA fluorescence. Populations were analyzed using FlowJo v.10.1.1 (FlowJo LLC, Ashland, OR).

### Extraction and sequencing of heterotroph DNA for genome assembly

Individual heterotrophs were grown in 45mL ProMM media in duplicate biological replicates and collected in late exponential growth. Cells were pelleted via centrifugation at 7,197 x g for 20 minutes at room temperature. The pellets were then extracted using a previously published standard phenol/chloroform procedure^44^, with modifications as previously described^22^. Genomic DNA was diluted and fragmented to 10-12kb using a gTube (cat# 520079, Covaris) and cleaned with 0.45x SPRIselect beads (cat# B23317, Beckman). Indexed SMRTbell libraries were prepared following the Template Prep kit 3.0 protocol (cat# 102-141-700, PacBio), pooled, and subjected to MinElute cleanup (cat# 102-194-100, PacBio) and sequenced on a PacBio Revio flow cell at the Harvard Bauer Core Facility.

### Reference genome assembly

Reference genomes were assembled using metaFlye v2.9.5 on PacBio long read sequencing with default parameters^45^. Taxonomic classification of assembled genomes was done using MMseqs2 v17.b804f with default parameters against the GTDB v207 database^46,47^. Genome quality was assessed using checkM2 v1.0.1, with all assembled genomes at >99% completeness and <1% contamination^48^. The protein sequences for the genomes were predicted using Prodigal v2.6.3^49^ and were annotated using eggnog-mapper v2.1.12^50^ and BlastKOALA^51^ using default settings (Supp. Table 10).

### Thalassospira strain comparison

Transcriptome experiments were conducted on *Thalassospira* strains prior to genome sequencing, with the aim of assessing potential strain- or phenotype-specific interactions with MED4. The two *Thalassospira* strains used had been isolated independently, so to determine their relatedness, we assembled their genomes via PacBio sequencing and compared these genomes using skANI v0.4.0^52^. The resulting ANI revealed the genomes were highly similar, at 99.99%. For additional checks, the genomes were aligned using MUMmer v3.23^53^ and 4 SNVs were detected, along with a large putative inversion of 2Mb^53^.

### Metagenomics sampling and extraction

To determine the composition of the heterotrophic community within our xenic MED4 culture, duplicate cultures were grown in 35mL Pro99 media each, combined and collected in late-exponential growth, and centrifuged at 7,197 xg for 25 minutes at room temperature. The pellet was then extracted using a previously published standard phenol/chloroform procedure^44^, with modifications as previously described^22^.

To determine the heterotroph composition in the synthetic community for each day of the growth curve, DNA was collected by filtering 5mL of the synthetic community cultures onto 0.2 μm 25 mm polycarbonate filters (Sterlitech, Auburn, WA). Filters were stored at -80°C until extraction. Prior to extraction, samples were thawed and internal standards (Thermus thermophilus HB27 DNA, ATCC BAA-163D-S) were added in proportion to the total amount of MED4 cells to serve as a control for DNA extraction efficiency. DNA was then extracted as previously described^24^.

### Metagenomics analysis

DNA libraries were prepared using a NexteraXT Index Kit v2 with a 1.5x SPRI cleanup before being sequenced at the MIT BioMicro Center on an Illumina NextSeq 500 Instrument to obtain 150bp paired-end reads. The genomic reads were trimmed using bbmap v39.01^54^ (bbduk -minlen 25 -trimq=10 maq=20 ktrim=r k=23 mink=11 hdist=1).

Reads from the xenic MED4 cultures were classified with kaiju v1.10.1^55^ using the kaiju_db_refseq_nr v2023-06-17 to obtain the relative read abundance of MED4 and heterotrophs.

The synthetic community metagenomic reads were mapped to the assembled reference genomes using coverm v0.7.0^56^ with bwa-mem v07.17^57^ (-m=mean --min-covered-fraction 0 --min-read-percent-identity 95). Internal standards were then corrected by the extraction efficiency of the *Thermus thermophilus* (ATCC BAA-163D-S) DNA, which had been added in ∼1% read abundance. Extraction efficiency was estimated using blastn v2.14.0 (-perc_identity 95). Genome equivalents of recovered *Thermus thermophilus* were then calculated using the formula:

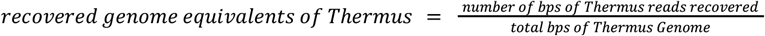

The recovered genomes were divided by the estimated number of genomes added, which was calculated using the following equations:

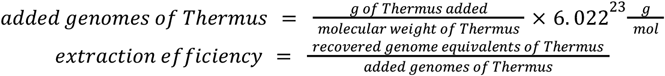

To ensure accurate comparisons within the community, we accounted for the estimated extraction efficiency of each individual microbe. Extraction efficiency for each heterotroph was calculated by dividing the estimated genome equivalents on Day 0 by the corresponding cell counts measured by flow cytometry on Day 0. Genome equivalents were determined from the number of mapped reads normalized by the total genome size of each organism. These Day 0 extraction efficiency values were then applied across the entire time course of the experiment (Supp. Table 3).

### RNA isolation and library preparation

Duplicate RNA samples were collected at Days 2, 4, and 5 by combining 12mL of culture with 36mL pre-chilled RNALater (4°C) and incubating at 4°C for 3-7days. Samples were then filtered onto 0.2μm 25mm polycarbonate filters (Sterlitech, Auburn, WA) and stored at –80°C. To quantify RNA per cell, internal standards were added in three 10uL pools prior to extraction. These standards were based on Gifford et al. (2016)^58^, with minor modifications to the PCR annealing temperatures to optimize on our equipment.

Total RNA was extracted from filters by first incubating each sample in 150μL10mM Tris (pH 8) with 1μL lysozyme (20,000 U, Thermo Fisher Scientific, Waltham, MA), and 2μL SUPERase-In RNase inhibitor (40 U, Invitrogen, Waltham, MA) for 5min at room temperature. RNA was then isolated using the mirVana miRNA extraction kit (Invitrogen, Waltham, MA) according to the manufacturer’s instructions. Extracted RNA was further purified with RNAClean XP Beads (Beckman Coulter, Brea, CA), resuspended in DEPC-treated water, and quantified using both Nanodrop (Thermo Fisher Scientific, Waltham, MA) and Ribogreen assays (Thermo Fisher Scientific, Waltham, MA). Ribosomal RNA was depleted as part of the KAPA HyperPrep with Ribo-Erase workflow. NEB Bacterial rRNA Depletion probes were substituted 1:1 from the probes included in the KAPA kit. cDNA synthesis, adapter ligation, and amplification were conducted following the KAPA workflow. After amplification, residual primers were eluted away using KAPA Pure Beads in a 0.63x SPRI-based cleanup. Pooled libraries were loaded onto an Illumina Novaseq6000 instrument to obtain either 100bp single-end or 111bp paired-end reads from the Harvard BPF Genomics Core Facility, and 50bp paired-end reads from the MIT BioMicro Center.

### RNAseq bioinformatics pre-processing and internal standard normalization

Reads were quality-filtered using bbduk v39.18 (minlen=25 qtrim=rl trimq=10 maq=20 ktrim=r k=23 mink=11 hdist=1) to remove low-quality regions and adapters^54^. Remaining reads were mapped to reference genomes and the internal standard sequences from the *S. solfataricus* P2 genome (IMG genome ID 638154518), using bowtie2 v2.5.4 with default parameters^59^. Read counts mapped to each feature were obtained using HTSeq v2.0.9 (-s reverse -t exon -r pos --nonunique all)^60^.

Quantitative transcriptomics counts were obtained following a modified version of the methods described previously^58^, utilizing synthesized RNA standards added at varying concentrations prior to sequencing. Sequencing efficiency was calculated from the number of recovered standard molecules using following the formula:

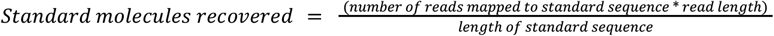

Linear regression was performed between the median number of standards recovered and the median number of standards added to estimate efficiency. Internal standards with median values outside three standard errors were removed (Supp. Fig. 10). The remaining internal standards were used to calculate efficiency rates (Supp. Table 7), which were then applied to normalize transcript counts:

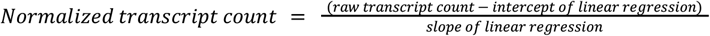

### RNA-seq bioinformatics processing

Internal standard-normalized transcript counts were summed across all sequencing runs (Supp. Table 7) to achieve sufficient depth, particularly for low-abundance heterotrophs. Significantly differentially expressed genes were identified using edgeR v4.2.1^61^, with p-values corrected for multiple hypothesis testing with the

Benjamini-Hochberg procedure. Genes with a q-value < 0.05 and log_2_-fold change >1.2 for MED4 or >1.5 for heterotrophs were considered significantly differentially expressed; the lower threshold for MED4 accounts for its weaker transcriptomic signal. Functional pathway enrichment of MED4 genes that were significantly differentially expressed in at least one condition were determined using clusterprofiler v4.12.6^62^ using the compareCluster function with enrichKEGG from BlastKOALA^51^. Gene expression clusters were obtained using euclidean clustering with the Pheatmap package.

To determine how each heterotroph responded to the community (relative to co-culture) the log_2_ fold change for each gene in each major KEGG pathway was averaged for each heterotroph. The KEGG pathways were determined using defined categories directly from KEGG BRITE^63^ and annotations from BlastKOALA. The mean log_2_ fold change was also determined for each KEGG sub-pathway, and results for the top sub-pathways in *Pseudohoeflea* and *Thalassospira* are shown in Figures 3B and 3C.

Transcript counts were corrected with flow cytometry data to obtain absolute transcripts per cell in the co-culture and synthetic communities for gene- and pathway-level analyses within the synthetic community. KEGG pathway analysis was performed using BlastKOALA^51^ See Code Availability section below for detailed subsequent analysis steps.

## Data Availability

All sequencing data associated with this manuscript are publicly available on NCBI under BioProject accession PRJNA1333906.

## Code Availability

All codes used to process and analyze metagenomics and metatranscriptomics sequencing data are available at our Github repository: https://github.com/jamesm224/community_dynamics

