## Supplementary Information for "The shared and distinct roles of *Prochlorococcus* and co-occurring heterotrophic bacteria in regulating community dynamics"

### Supplementary Figures

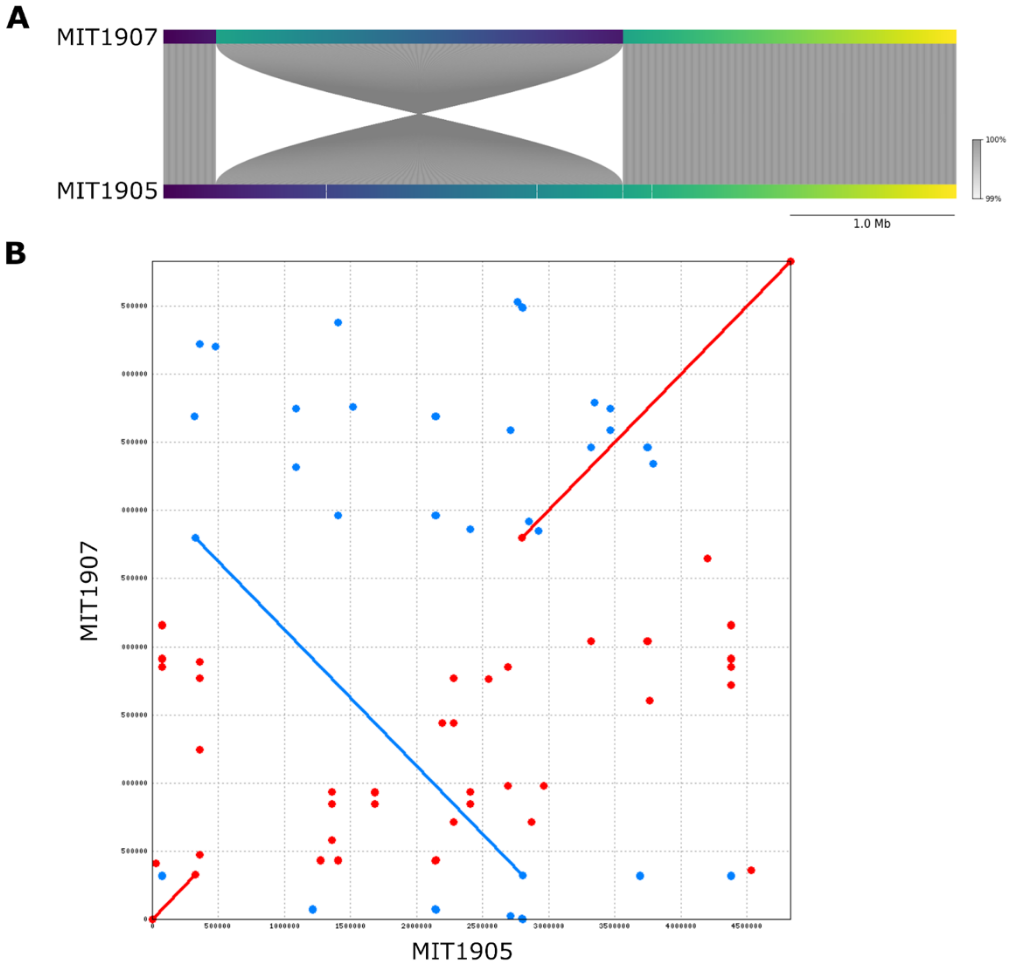


**Supplementary Figure 1: Comparison of *Thalassospira* genomes. A)** Genome plot obtained using FastANI, comparing the two isolated *Thalassospira* strains. Four SNVs and one large (~2Mb) inversion are seen. **B)** Dot plot of the genome alignment of the two isolated Thalassospira strains using MUMmer. The blue line indicates the same large inversion seen in A.

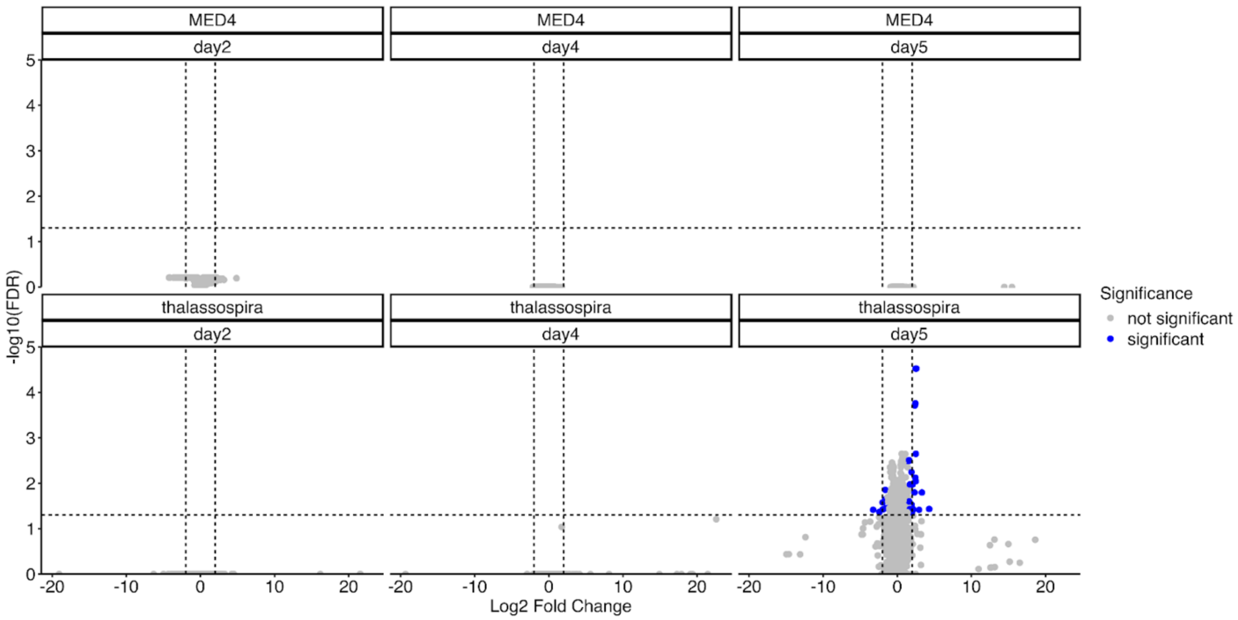


**Supplementary Figure 2: Comparison of the transcriptomic responses of the two *Thalassospira* strains.** Each column represents a day (2, 4, or 5) of the growth curve. Top row indicates log2 fold changes in MED4 gene expression when co-cultured with MIT1905 vs MIT1907. No significantly differentially expressed genes are observed for any day in MED4. Bottom row indicates log2 fold changes in Thalassospira MIT1905 vs MIT1907 gene expression when co-cultured with MED4. No significant differentially expressed genes were observed for Days 2 and 4, but some are observed for Day 5 (blue dots), likely due to noise in the cultures. Y-axis shows the log10 q-values.

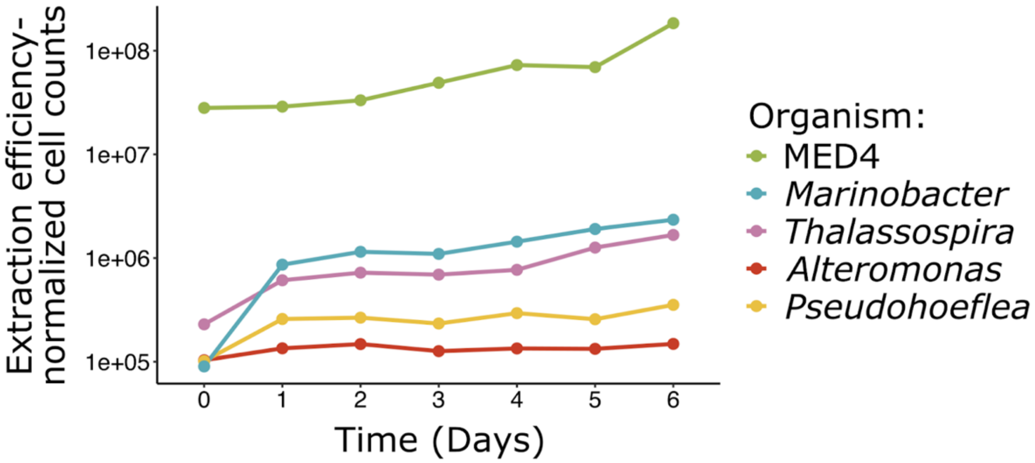


**Supplementary Figure 3: Community composition across the MED4 growth curve.** Estimated cell counts for each day of the MED4 growth curve in the synthetic community. Metagenomic sequencing results were normalized by internal standards and by extraction efficiency to determine estimated cell counts. Colors indicate each organism in the community.

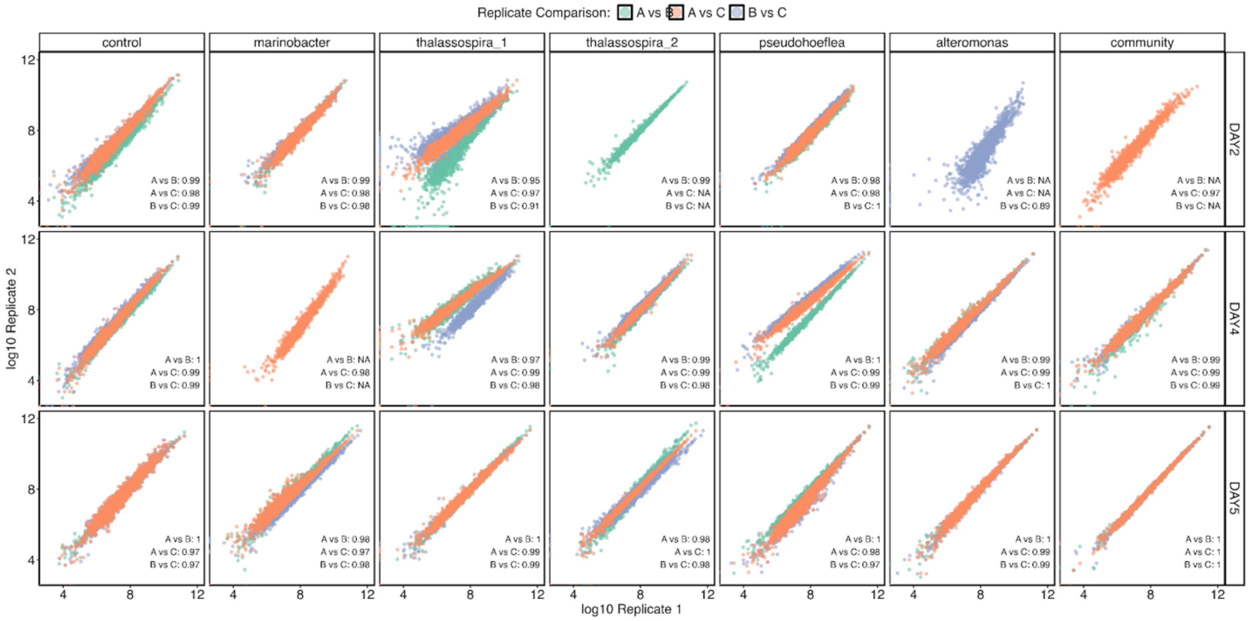


**Supplementary Figure 4: RNA-seq replicate comparison plots.** Comparison of RNA-seq across all three biological replicates for each culture condition and day. Rep A vs B (green), Rep A vs C (orange), and Rep B vs C (blue). R-squared correlations are listed for each sample comparison.

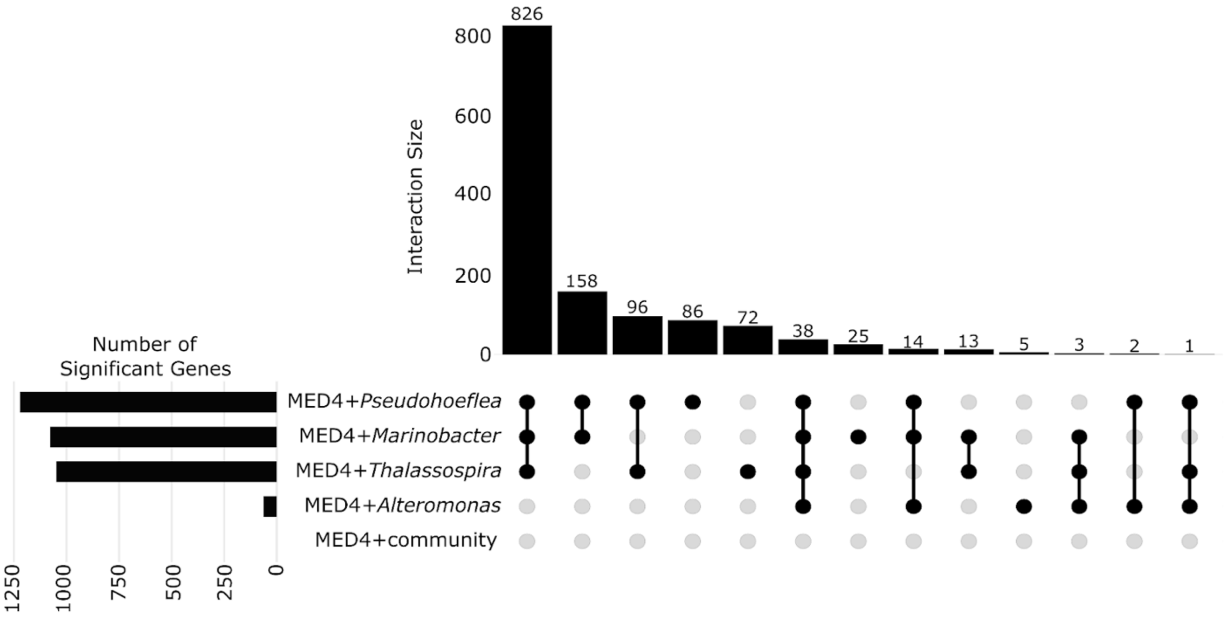


**Supplementary Figure 5: Quantity and relationship of significantly differentially expressed MED4 genes on Day 2.** Left, bar plot of the total number of significantly differentially expressed MED4 genes in each culture condition relative to axenic MED4 (control). Right, number of overlapping significantly differentially expressed genes across conditions (interaction size). Black circles and lines indicate which conditions are being compared.

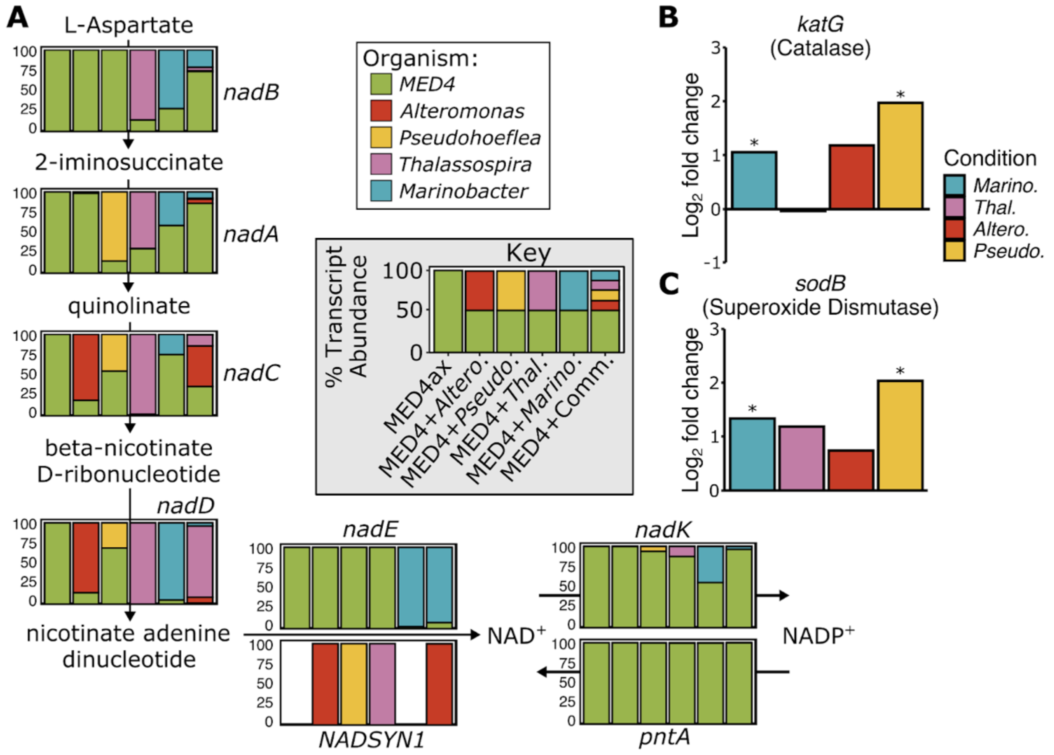


**Supplementary Figure 6: Production of communal NAD$^+$, catalase, and superoxide dismutase in each condition. A)** Metabolic pathway for NAD$^+$ biosynthesis showing the relative abundance of each transcript coming from each organism for each gene in the pathway. **B)** $Log_2$ fold change of *katG* (catalase) abundance in each heterotroph grown in community relative to co-culture. Positive values indicate the transcript is enriched in the community relative to co-culture. * indicates q<0.05. **C)** Same as **B,** except for *sodB* (superoxide dismutase)

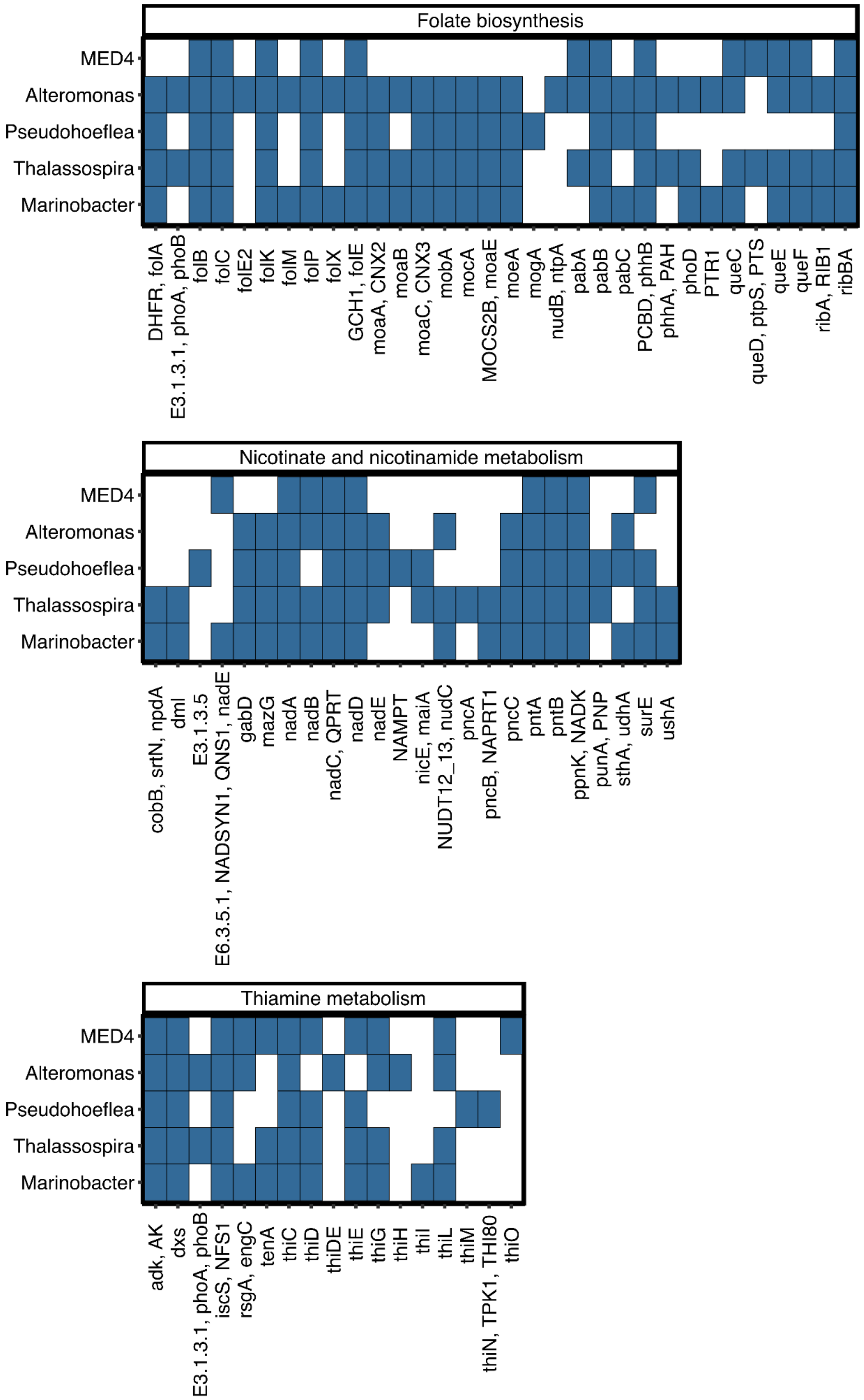


**Supplementary Figure 7: Presence/absence of genes in relevant B vitamin biosynthetic pathways.** Genes on the x-axis involved in folate biosynthesis (top), nicotinate and nicotinamide metabolism (middle), and thiamine metabolism (bottom). Y-axis indicates organism. Blue squares indicate the gene is present, white squares indicate absence.

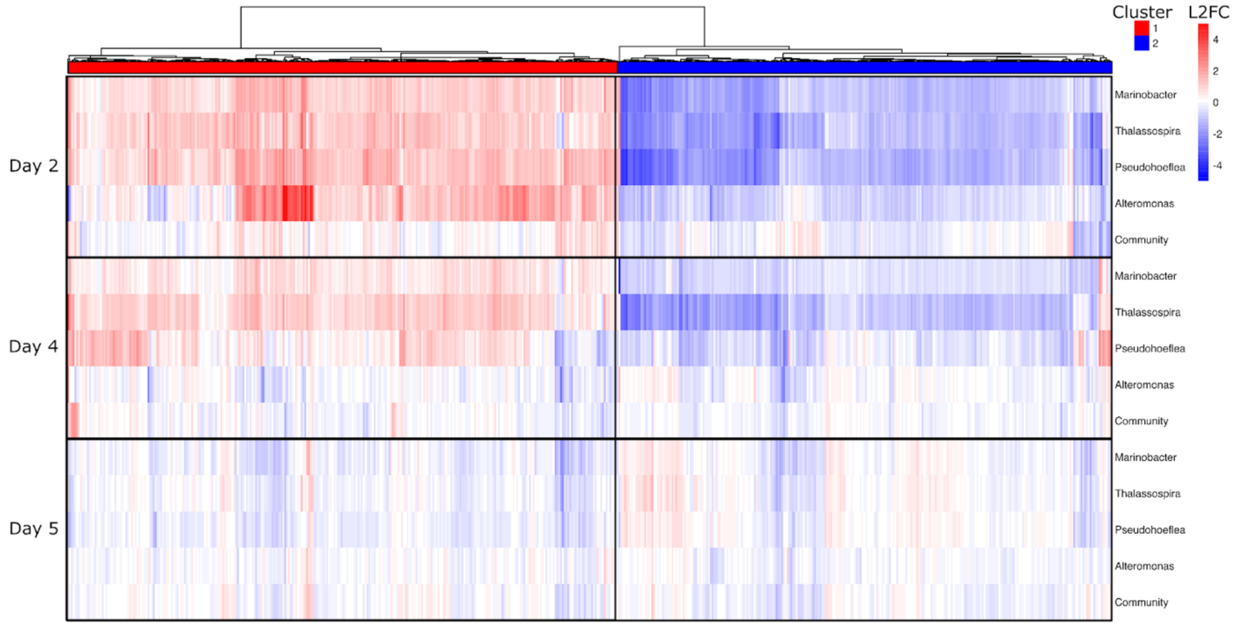


**Supplementary Figure 8: Effects of time on MED4 gene expression.** Heatmap of the $\log_2$ fold change (LFC) for all MED4 genes significantly differentially expressed in at least one condition relative to growth alone. Genes are clustered on the x-axis by similarity across treatments. Negative values in blue indicate higher expression in axenic MED4, and positive (red) values indicate higher expression in MED4 grown with heterotrophs. Y-axis indicates the condition, which is broken down into Day 2 (top), Day 4 (middle), and Day 5 (bottom).

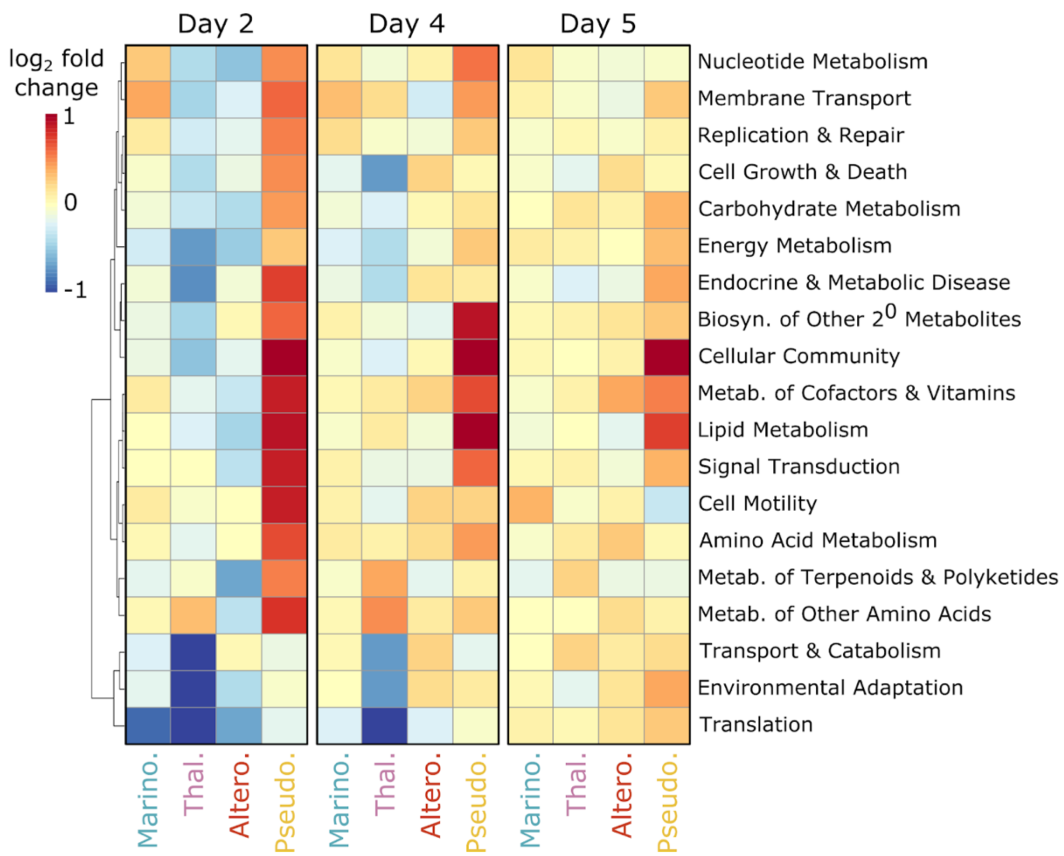


**Supplementary Figure 9: Effects of time on heterotroph gene expression.** Heatmap of the mean $\log_2$ fold change in transcript abundance for each major KEGG pathway for each heterotroph when growth in individual co-culture versus its growth in the synthetic community. Positive (red) values indicate higher expression in the co-culture relative to the community. Values are shown for Days 2 (left), 4 (middle), and 5 (right).

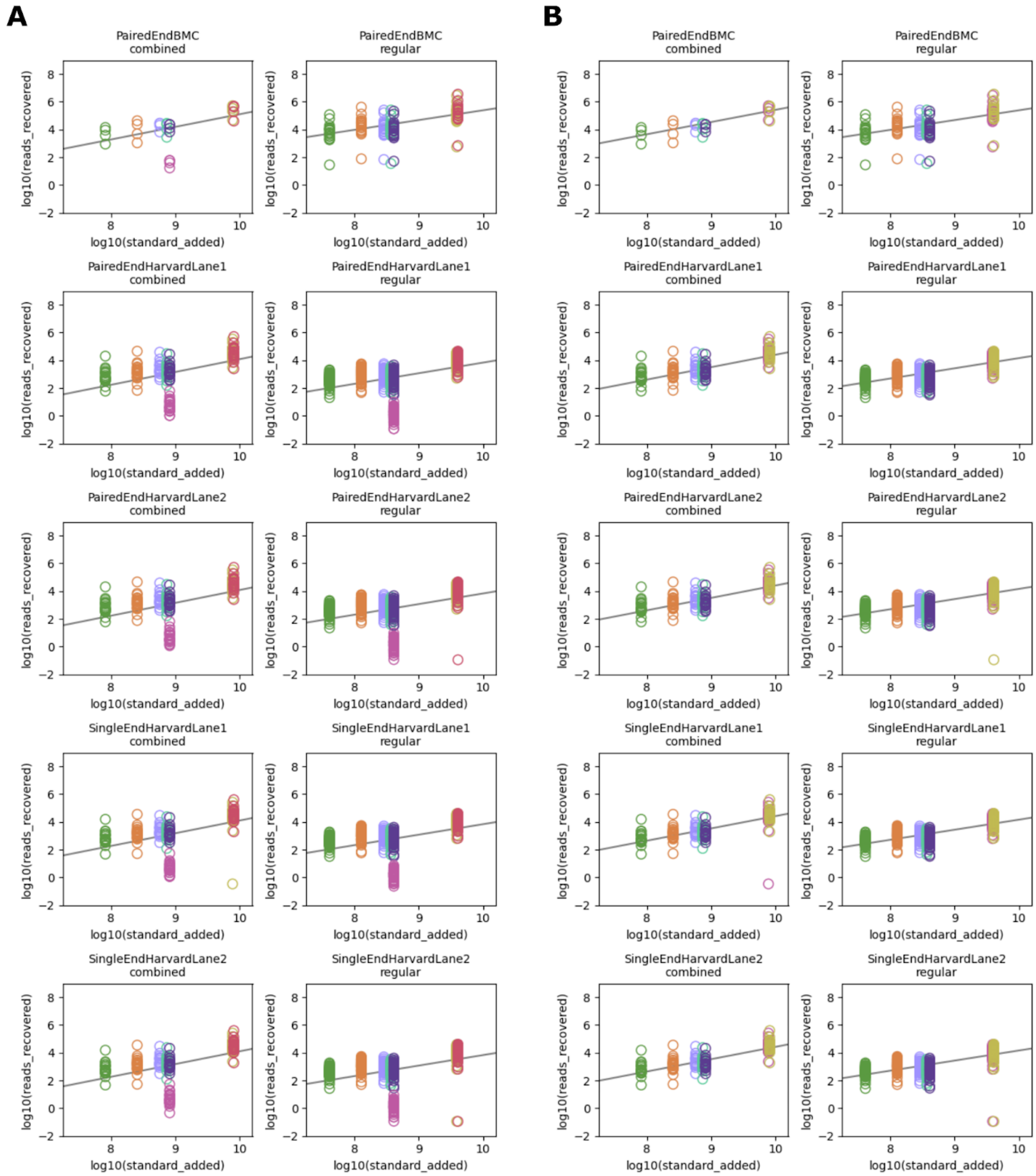


**Supplementary Figure 10: RNA-seq internal standard normalization plots.** Linear regression for all RNA-seq internal standards comparing the amount of standard added vs the amount of standard recovered. Each plot represents a different sequencing run and lane. **A)** Pre-filtered internal standards. **B)** Post-filtering to remove and standards outside 3 standard errors.

### Supplementary Tables:

**Supplementary Table 1: Strains used in this study**

**Supplementary Table 2: Flow cytometry counts**

**Supplementary Table 3: Absolute metagenomic count data - including extraction efficiency, genome recovery, and genome equivalents**

**Supplementary Table 4: ANI genome comparison between the two *Thalassospira* strains and between previously sequenced Thalassospira strains**

**Supplementary Table 5: *Thalassospira* RNA-seq comparison**

**Supplementary Table 6: MED4 significantly differentially expressed genes in each condition**

**Supplementary Table 7: Internal Standard Normalization**

**Supplementary Table 8: *Marinobacter*, *Thalassospira, Alteromonas*, and *Pseudohoeflea* significantly differentially expressed genes**

**Supplementary Table 9: Absolute transcript counts**

**Supplementary Table 10: Genome annotations**
